# Coilin oligomerization and remodeling by Nopp140 reveals a hybrid assembly mechanism for Cajal bodies

**DOI:** 10.1101/2021.08.01.454609

**Authors:** Edward Courchaine, Martin Machyna, Korinna Straube, Sarah Sauyet, Jade Enright, Karla M. Neugebauer

## Abstract

Cajal bodies (CBs) are ubiquitous nuclear membraneless organelles (MLOs) that promote efficient biogenesis of RNA-protein complexes. Depletion of the CB scaffolding protein coilin is lethal for vertebrate embryogenesis, making CBs a strong model for understanding the structure and function of MLOs. Although it is assumed that CBs form through biomolecular condensation, the biochemical and biophysical principles that govern CB dynamics have eluded study. Here, we identify features of the coilin protein that drive CB assembly and shape. Focusing on coilin’s N-terminal domain (NTD), we discovered its unexpected capacity for oligomerization *in vivo*. Single amino acid mutational analysis of coilin revealed distinct molecular interactions required for oligomerization and binding to the Nopp140 ligand, which facilitates CB assembly. We demonstrate that the intrinsically disordered regions of Nopp140 have substantial condensation properties and suggest that Nopp140 binding thereby remodels stable coilin oligomers to form a particle that recruits other functional components.

## Introduction

Cajal bodies (CBs) are membraneless organelles (MLOs) present in the nuclei of most vertebrate, insect, and plant cells with a catalog of molecular components^1,2^. Discovered by Ramon y Cajal in 1903, CBs are an important model for understanding the assembly, maintenance and function of MLOs due to extensive knowledge about their composition and function. CBs are the sites of small non-coding ribonucleoprotein particle (RNP) assembly and recycling. Nucleolar snoRNAs and spliceosomal snRNAs traffic through CBs for their modification and maturation to functional components of the splicing and ribosomal processing machinery, namely snRNPs and snoRNPs^3–7^. CBs are also nucleated at specific sets of genes, such as those encoding snRNAs, thereby clustering whole chromosomal regions within the three-dimensional space of the nucleus^3,8–10^. Like other MLOs, CBs are dynamic and contain numerous molecular constituents that freely diffuse in and out of CBs with residence times of less than 1 minute^5,11,12^. They disassemble before mitosis and assemble thereafter, are absent in G0, and are modified to respond to cellular and organismal stress^13^.

Unlike many other MLOs, at least one important function of CBs themselves is known. The ability to deplete or genetically delete the acknowledged “scaffolding factor” of CBs, coilin, provides a handle for identifying deficits upon CB loss in plants and animals *in vivo*^14–18^. Although phenotypes differ among species, coilin is essential for vertebrate embryonic survival and for mammalian fertility^14,15,19–21^. Loss of viability due to inefficient snRNP assembly and splicing supports evidence that the rate of snRNP assembly is faster in CBs, allowing the biogenesis of components of the splicing machinery to keep pace with zygotic gene expression during very short cell cycles^14,22,23^. Moreover, CB composition and morphology are significantly altered in human disease, such as Spinal Muscular Atrophy (SMA) and Down Syndrome^24–26^.

Despite this wealth of information and their obvious physiological importance, the field lacks a mechanistic understanding of how CBs assemble. Because of the absolute requirement for coilin, the answer must lie with its structure and function. Coilin contains three regions that are separable by the degree of evolutionary conservation^2^. The N-and C-terminal domains (NTD and CTD, respectively) are the most highly conserved; they are separated by a region of low complexity that is poorly conserved and likely intrinsically disordered. This middle domain contains a bipartite nuclear localization signal (NLS) and an RG box modified by dimethylarginine and bound by the spinal motor neuron protein (SMN) tudor domain^11,27–30^. The coilin CTD harbors its own tudor domain, which interestingly lacks the aromatic amino acids that would enable it to bind dimethylarginine ^31^. Instead, the coilin CTD binds to the core Sm ring of spliceosomal snRNPs that are defining components of CBs^28,32,33^. The coilin CTD likely binds several other proteins and determines the number of CBs per cell^2,34,35^.

The coilin NTD is required for CB formation and for targeting coilin to existing CBs: the NTD alone can localize to CBs formed by full-length coilin and, conversely, coilin constructs lacking the NTD cannot^35^. Because the NTD is also required for the pull-down of endogenous coilin from cell extracts by tagged and expressed coilin constructs, the NTD has been referred to as a “self-interaction” domain. Indeed, coilin does self-associate both in the nucleoplasm and in CBs, as detected by FRET^4^. However, it has never been clear whether coilin NTD binds directly to the NTDs of other coilin molecules, if it forms higher-order oligomers, or if the NTD binds to additional constituents that mediate CB assembly.

Here, we dissect the molecular functions of the coilin NTD *in vivo*, combining an alanine scanning screen of conserved single amino acids with the expression of protein constructs in either the nucleus or cytoplasm, in cells containing endogenous coilin and CBs, and in cells lacking endogenous coilin. These experiments sample the ability of the coilin NTD to self-associate in different cellular contexts, facilitating the discovery that the domain is capable of extensive oligomerization and identifying Nopp140 as the critical nuclear ligand for CB formation. Finally, we used protein structure prediction to visualize the positions of the amino acids critical for oligomerization and Nopp140 binding, allowing us to propose a working model for how the interplay between these two activities contribute to CB assembly.

## Results

### The Coilin NTD undergoes extensive homo-oligomerization in the cytoplasm and forms nuclear puncta containing Nopp140

To investigate the hypothesis that the coilin NTD promotes CB assembly through dimerization activity, we asked if the NTD could be replaced by a heterologous protein dimerization domain. Thus, we cloned a set of expression constructs (Fig. 1a) comprised of full-length coilin, coilin deleted of the 97 amino acid NTD, and coilin having the NTD replaced by the leucine zipper from GCN4^36^. Each construct was transfected into mouse embryonic fibroblasts derived from coilin gene knock-out mouse (*Coil-/-* MEFs), which thereby lack CBs^15^. Each protein was well-expressed (Fig. S1a). Fluorescence resonance energy transfer (FRET) assays detected significant interactions between full-length coilin molecules (coilin^FL^-coilin^FL^), as shown previously ^4^, and between those dimerized by the GCN4 leucine zipper (coilin^GCN4^-coilin^GCN4^) confirming the functionality of the GCN4 dimerization domain (Fig. S1b). As expected, coilin^FL^ induced the formation of CBs in *Coil-/-* MEF nuclei (Fig. 1b), containing typical snRNP components of CBs (Fig. S1c). Coilin^ΔNTD^ did not support intramolecular interaction as assayed by FRET (Fig. S1b) and was unable to form CBs, as previously reported^35^. It was, therefore, surprising that dimerization by coilin^GCN4^ did not change the punctate distribution of fluorescence observed for coilin^ΔNTD^. Both sets of puncta – a term we use throughout the manuscript to signify clusters of fluorescence – are smaller and more numerous than typical CBs, leading us to conclude that dimerization by the N-terminus of coilin is not sufficient for proper CB formation. Rather, the coilin NTD may facilitate CB formation through specific activities in addition to simple dimerization.

**Fig 1.**
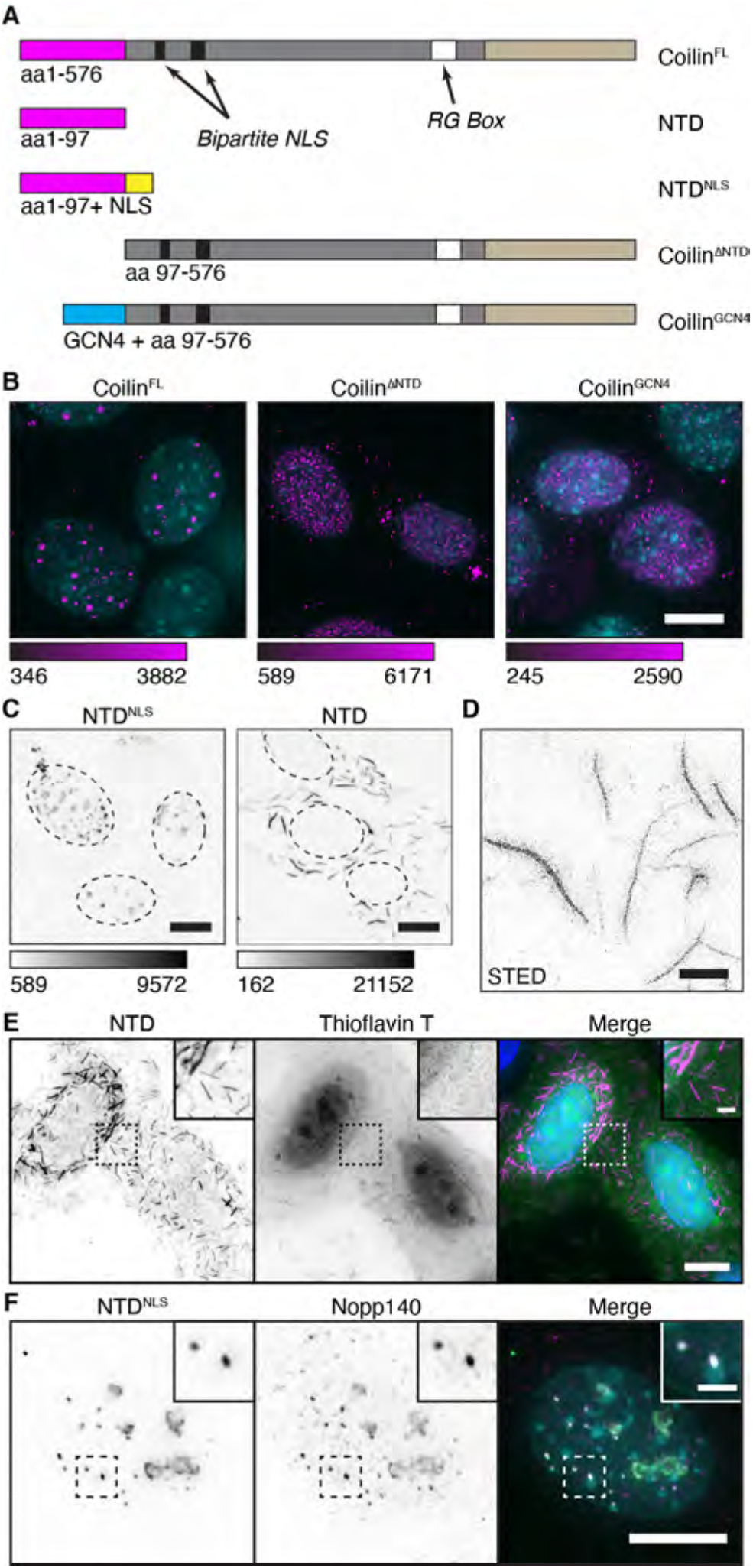
The coilin NTD assembles into oligomers or puncta *in vivo*. A) Linear schematics of coilin constructs used in this study. The number of amino acids (576 aa in full-length human coilin) included in the expressed protein is indicated below and to the left of each diagram. Coilin NTD is pink, the C-terminus is beige, the intrinsically disordered central region is shown in gray, the NLS shown in yellow is from SV40, and the GCN4 leucine zipper dimerization motif is in cyan. The NTD and NTD^NLS^ constructs were tagged with either myc- or HA-tags on their C-termini (not shown). B) *Coil^−/−^* MEFs transfected with coilin^FL^, coilin^ΔNTD^ (a deletion construct lacking the NTD) or coilin^GCN4^ with attempted complementation of the NTD by the heterologous dimerization motif. Color bars are in analog-digital units to indicate differences in expression. C) Grayscale images of *Coil^−/−^* MEFs transfected with an expression construct encoding coilin NTD, lacking an NLS. Confocal images with scale bar 10 μm. Dotted lines indicate the limits of each nucleus. D) STED; scalebar 2 μm. E) Cytoplasmic NTD oligomers in HeLa do not stain with Thioflavin T. F) *Coil^−/−^* MEF transfected with NTD^NLS^ and counterstained for Nopp140. All images acquired with DeltaVision except C and D. Scale bars = 10 μm; insets 2 μm.

To investigate the independent function(s) of the coilin NTD isolated from the rest of the coilin molecule, which has numerous binding partners^2^, we expressed the coilin NTD in the cytoplasm or in the nucleus by utilizing a viral SV40 nuclear localization signal (Fig. 1a). Unexpectedly, the NTD construct lacking the NLS formed abundant, very long oligomers up to 10 micrometers long in the cytoplasm (Fig. 1c). Despite this length, the oligomers appear to be as little as 100 nm in diameter by sub-diffraction imaging (Fig. 1d). Using aggregation-sensitive dyes (Figs 1e and S1c), we showed that these cytoplasmic NTD oligomers do not appear to be amyloid in nature; Thioflavin T and Congo Red typically recognize amyloid, while ProteoStat labels aggregates. In contrast to this cytoplasmic behavior, expression of the NTD^NLS^ protein in the cell nucleus led to the formation of apparently round MLOs that appeared to be similar in size and number to CBs (see Figs 1c and 1f). However, the nuclear puncta formed by NTD^NLS^ did not contain the usual snRNP or snoRNP components of CBs marked by SART3, SmB”, or fibrillarin (Fig. S1d), indicating the large NTD^NLS^ puncta are not *bona fide* CBs. A notable exception was Nopp140, a nucleolar and CB protein that has previously been shown to bind the coilin NTD^37,38^ and has been linked with the accumulation of snoRNP components in CBs^6^. We conclude that the NTD has the capacity to assemble into extensive oligomers or, alternatively, into large puncta depending on nuclear or cytoplasmic localization.

The observed association of Nopp140 with nuclear NTD^NLS^ in puncta raised the question of whether Nopp140 binding to the coilin NTD contributes to CB assembly, which is a controversial point in the literature. Although one previous publication reported that CRISPR targeting of both alleles of Nopp140 did not disturb CB formation^6^, two other studies showed that loss of Nopp140 is correlated with CB disassembly^38,39^. To determine the requirement of Nopp140 for CB assembly and maintenance in our hands, we depleted Nopp140 protein from HeLa cells using siRNAs (Fig. 2A; note that the NOLC1 gene encodes Nopp140 protein). Untreated HeLa cells contain ~2 coilin-positive CBs per nucleus and slightly fewer SMN-containing MLOs, called gems. Gems lack coilin and do not overlap CBs, although they can be closely associated with CBs^30,40^. As a positive control, coilin depletion reduced the number of CBs to less than half the original number per nucleus without affecting the number of gems, as expected (Figs 2b–e). Remaining CBs marked by coilin reflect variability in the efficiency of transfection (see Figs 2b&e). Importantly, depletion of Nopp140 significantly reduced the number of CBs (Fig. 2b) with a noticeable increase in the intensity of coilin staining outside of CBs (see Fig. 2d–f, line traces). Taken together, these data indicate that Nopp140 contributes to the localization of nucleoplasmic coilin in CBs.

**Fig 2.**
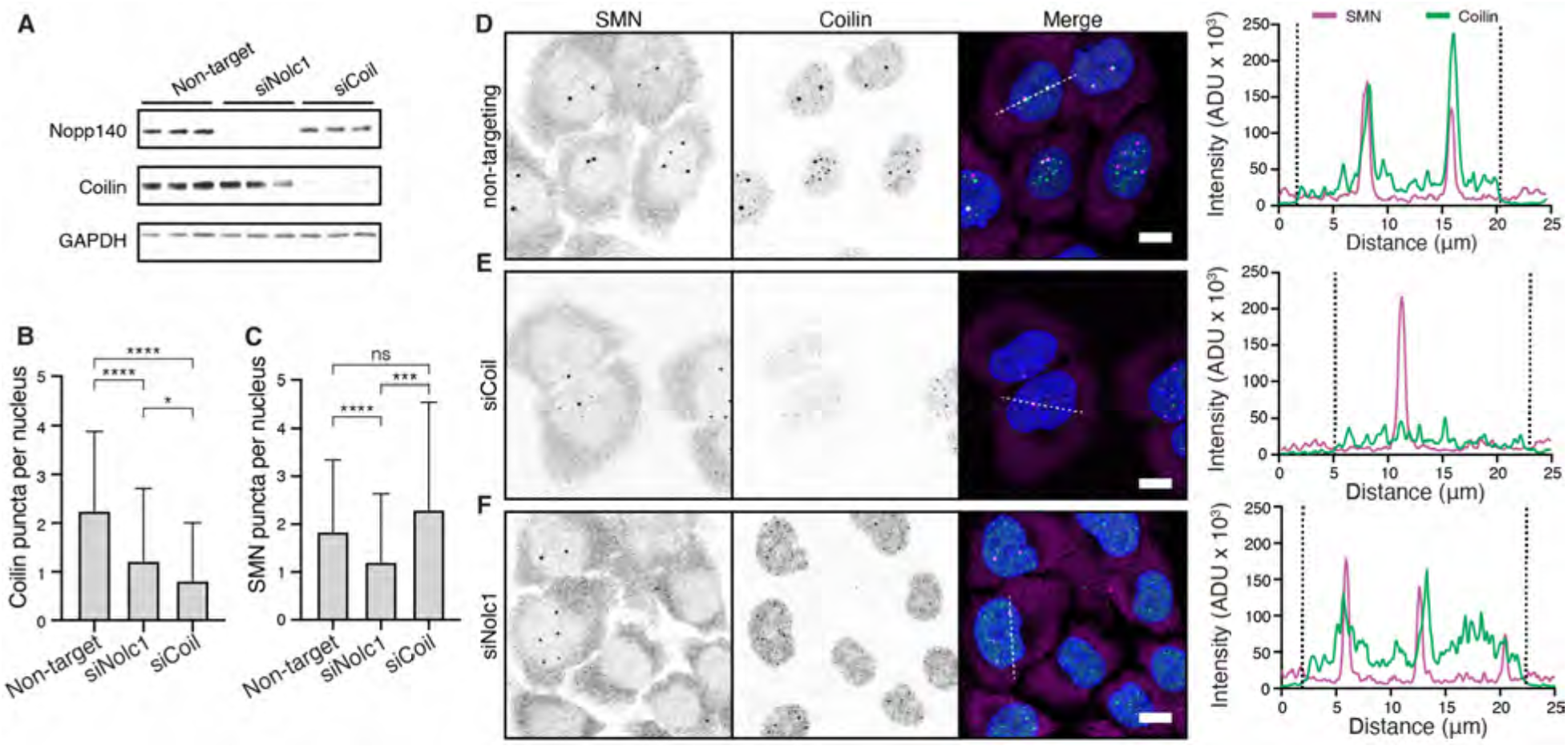
Nopp140 is essential for proper Cajal body assembly. A) Western blots of total lysate from three replicates of HeLa cells depleted of Nopp140 or coilin using siRNA transfection targeting their transcripts *Nolc1* and *coil*, respectively. B&C) Bar plots indicating the number of coilin puncta (B) or SMN puncta (C) per nucleus in cells undergoing depletion. Error bars indicate standard deviation. For each condition, 30-50 cells were counted for each of three biological replicates and then pooled. (*) indicates p < 0.05, (***) indicates p < 0.001, and (****) indicates p < 0.0001 for Mann Whitney test. D-F) Representative images and corresponding line profiles of summed projections for cells undergoing transfection with a non-targeting oligo (D), si*Coil* oligo pool (E), or si*Nolc1* oligo pool (F). Scale bars = 10 μm. Dotted lines in line profiles indicate the nuclear boundaries. Images acquired with Leica Sp8 laser scanning confocal microscope.

### Single amino acid mutations in the coilin NTD reveal discrete coilin-coilin and coilin-Nopp140 interactions

To gain mechanistic insight into how the coilin NTD forms oligomers in the cytoplasm and MLOs with Nopp140 in the nucleus, we performed an alanine scan by individually replacing each of 29 highly conserved amino acids present in the NTD (red shading in Fig. 3a) in the context of 29 individual full-length coilin expression constructs that were each introduced into *Coil-/-* MEF cells that lack CBs. These amino acids are more broadly conserved throughout evolution, and many occur within predicted beta sheet and alpha helix motifs defined as a ubiquitin-like fold according to RaptorX (Fig. S2). Figure 3b shows three single amino acid mutants – R8A, R36A, and D79A – that led to noticeable differences in the MLOs formed by coilin^FL^ compared to wild-type (WT), which formed large, discrete CBs with little observable fluorescence in the surrounding nucleoplasm (see also Fig. 1b). By comparison, R8A and D79A yielded small puncta and hazy nucleoplasmic background; R36A was chosen for contrast, because it had a subtler phenotype. For representative images of all remaining single amino acid mutations, showing a range of phenotypes, see Fig. S3.

**Fig 3.**
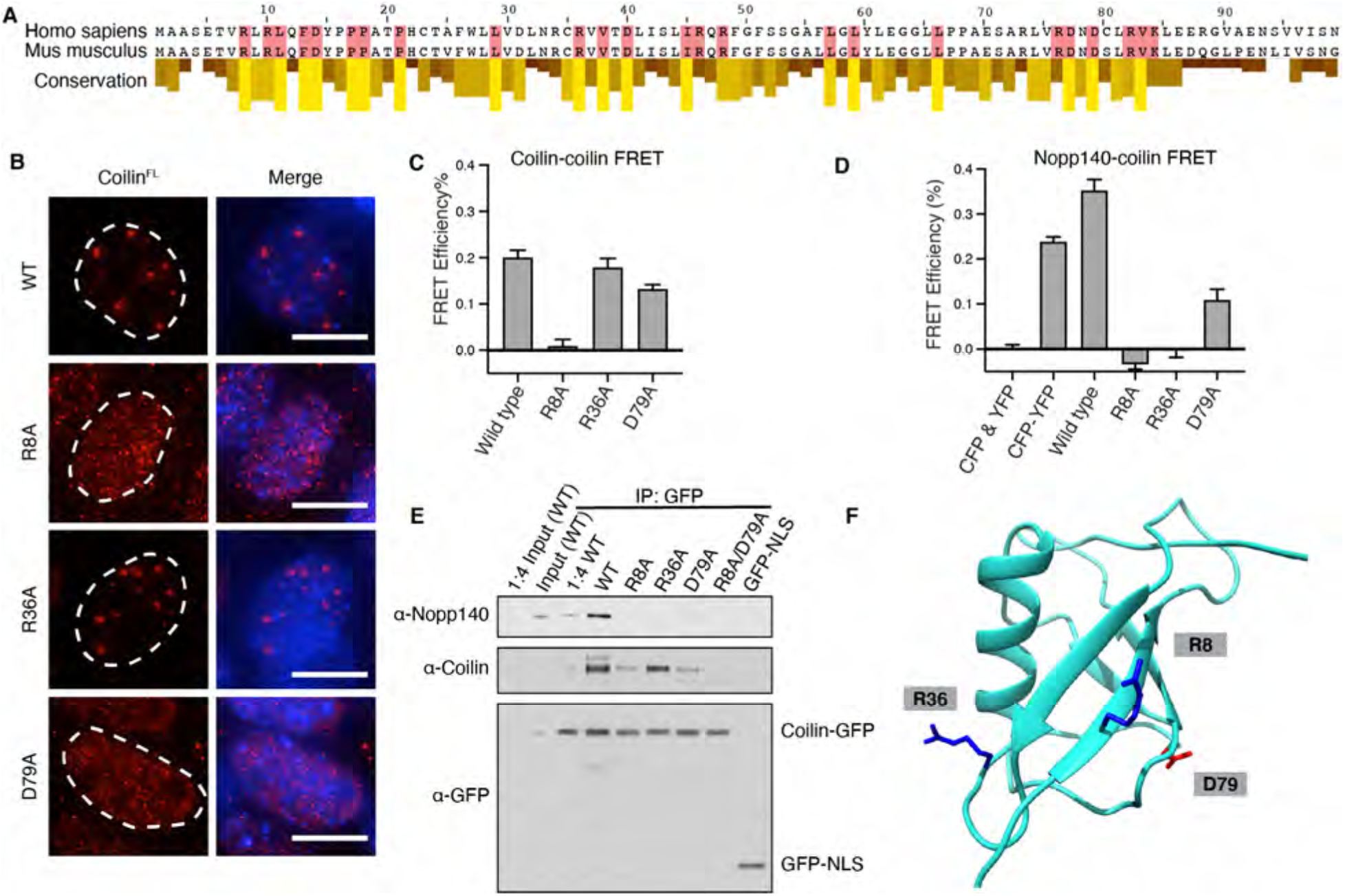
Point mutations in the coilin NTD affect self-interaction and Nopp140 association. A) Sequence alignment between mouse and human coilin NTDs with conservation scores taken from a Clustal Omega multiple alignment of vertebrate coilin sequences. Residues highlighted in red have high conservation and were selected for mutagenesis to alanine. B) Transfection of wild-type or different mutations of coilin^FL^ were transfected into *Coil-/-* MEF cells and examined for CB formation as a visual assay (coilin visualized in red, DNA visualized in blue). Scale bars = 10 μm. C&D) *In vivo* FRET measurements using the acceptor photobleaching method wherein apparent FRET efficiencies were calculated. Images acquired with DeltaVision. C) between coilin-CFP as the donor molecule and coilin-YFP as the acceptor co-expressed in HeLa cells and D) between coilin-CFP and Nopp140-YFP. The indicated amino acid changes were made in all coilin molecules tested, where indicated. Error bars represent standard error of the mean. E) Co-immunoprecipitation performed from HeLa cell total lysate after transfection with wild-type and mutant coilin-GFP constructs, as indicated. Coilin-GFP acts as the bait for either endogenous coilin or Nopp140 as prey and detected by Western blotting as indicated. F) Structure prediction of human coilin NTD aa1-97, which should adopt a ubiquitin-like fold. Two arginines and one aspartate implicated in coilin oligomerization and binding to Nopp140 are indicated.

If any of the single amino acid mutations in the NTD disrupted coilin-coilin and/or coilin-Nopp140 interactions, as intended, our acceptor photobleaching FRET assay (see Fig. S1b) would detect these changes. Wild-type coilin-coilin FRET yielded ~20% apparent FRET while wild-type coilin-Nopp140 FRET yielded ~35% (Figs 3c&d). Interestingly, the R8A mutation nearly abolished coilin-coilin FRET and D79A reduced it. This loss of FRET suggests the interaction could be direct. Our presumption that some of the single amino acid hits could perturb Nopp140-coilin interactions was borne out as well, because R8A and R36A abolished this signal, while D79A was again reduced.

To validate these interactions detected by FRET, pull downs were conducted from cellular lysates expressing GFP-tagged coilin. Wild-type coilin-GFP efficiently pulled down endogenous coilin and Nopp140 present in the extract (Fig 4e). In agreement with the FRET measurements, R8A and D79A mutations in the expressed, GFP-tagged coilin reduced the amount of endogenous coilin pulled down. In contrast, coilin-R36A-GFP pulled down normal amounts of endogenous coilin; yet, coilin-GFP constructs harboring any of the three mutations failed to pull down Nopp140. We predicted three-dimensional structure of the coilin NTD using RaptorX ^41^ and inspected the positions of R8, R36 and D79 (Fig. 4f), which are all required for CB assembly and maintenance (Figs 3 and S3). Taking the pulldown data together with the FRET data, the predicted structure suggests that R36 is accessible to Nopp140 on one face of the NTD, while R8 and D79 are accessible to other coilin monomers on the opposite face. Because R8A affects both coilin-coilin and Nopp140-coilin interactions in the FRET and pulldown assays, we speculate that Nopp140 may require coilin-coilin oligomerization for optimal binding.

**Fig 4.**
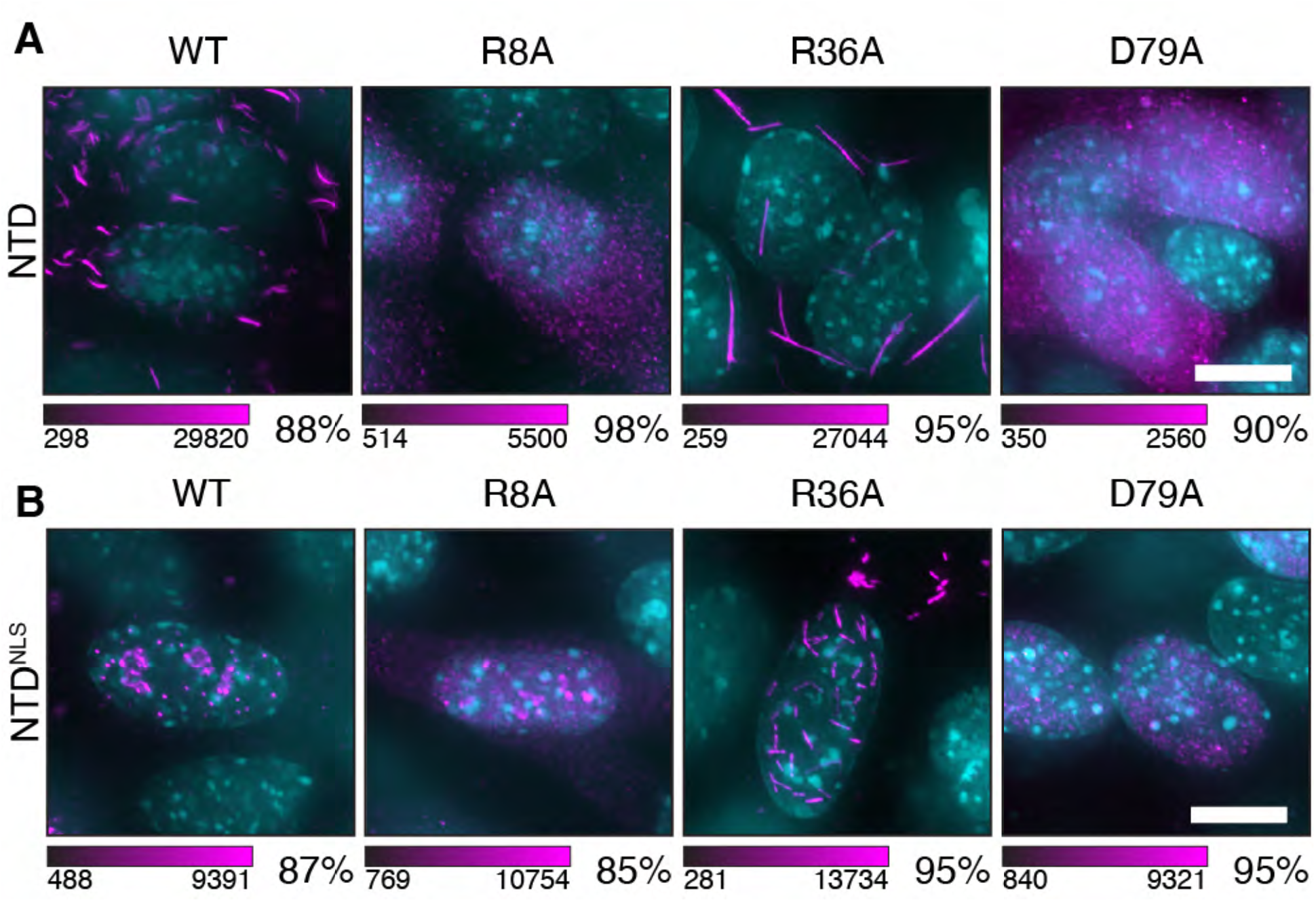
Point mutations alter the NTD’s capacity to oligomerize and form puncta. *Coil^−/−^* MEFs transfected with constructs for cytoplasmic expression (A, NTD) or for nuclear expression (B, NTD^NLS^) of the coilin NTD bearing selected point mutations. The NTD was immunostained with anti-myc (magenta) and DNA visualized with DAPI (cyan). Color scale bars given in analog-digital units. Scale bars = 10 μm. Percentage to the lower right indicates the fraction of cells with the displayed phenotype where n = 300 cells. Images acquired with DeltaVision.

### Nuclear Nopp140 modulates coilin oligomerization

To test the capacity of the NTD to oligomerize and form puncta in relation to the proposed self- and Nopp140 interactions, we utilized the single amino acid mutations in the context of the NTD expression constructs alone. Wild-type and mutant NTD or NTD^NLS^ were transfected into *Coil-/-*MEFs cells, as done previously (see Fig 1a). Wild-type NTD was localized to the cytoplasm and formed extensive oligomers as expected (Fig. 4A). However, R8A and D79A mutations abolished oligomer formation while the R36A mutation did not, thus confirming the results of our screen and FRET experiments. Note that the previous FRET measurements were also consistent in the context of the NTD alone (Fig. S4A); wild-type NTD-NTD FRET signals approached 40% and were sensitive to R8A and D79A mutations, while R36A had no effect on NTD-NTD FRET. In the nucleus, wild-type NTD^NLS^ formed puncta and some localization to nucleoli was observed (Fig. 4B). In this case, the R8A mutation led to the loss of nucleoplasmic puncta with associated nucleolar protein still present; the D79A mutation more strongly disrupted all puncta owing to the loss of coilin-coilin interactions (Fig. S4A). Remarkably, the R36A mutation, which effects Nopp140-coilin but not coilin-coilin interaction, caused oligomerization of NTD^NLS^ in the nucleus (Fig. 4B)! This finding is in strong agreement with our proposal that the R36A mutation disrupts Nopp140 binding, providing an explanation for why the wild-type NTD forms oligomers only in the cytoplasm: Nopp140 is a strictly nuclear protein. Moreover, the observation that R8A and D79A are essential for both oligomerization and the formation of puncta also agrees with our proposal that these amino acids mediate coilin-coilin interactions.

### Single amino acid coilin mutations have dominant negative effects on endogenous CBs, implicating homo-oligomerization in CB assembly

The dramatic effects of the single amino acid mutations on the formation of coilin assemblies in *Coil-/-* MEF cells – in the absence of endogenous coilin – led us to wonder how these mutants would interact with the structure of pre-existing CBs, i.e. in the presence of endogenous coilin. To test this, we expressed the NTD and NTD^NLS^ constructs in normal HeLa cells that express wild-type coilin and have CBs (see Fig. 2). As shown in Figure 5 the wild-type NTD forms oligomers in the cytoplasm of HeLa cells similar to those we observed in mouse cells, while the NTD^NLS^ protein readily enters and concentrates in CBs and nucleoli. Quantification of the number of CBs per nucleus reveals that neither wild-type NTD construct significantly alters CB number (Figs 5a and 5c). In sharp contrast, all NTD mutants – R8A, R36A and D79A – essentially eliminate round, nuclear CBs in transfected cells. This dominant negative disassembly of all CBs by R8A and D79A is apparently due to the remaining contacts with wild-type coilin (see Fig. 3e). Remarkably, the gain-of-function effects of R36A on CBs are due to the recruitment of endogenous coilin molecules by both NTD and NTD^NLS^ oligomers, evidently depleting coilin from the nucleoplasm and CBs. The data substantiate the notion that R8 and D79 are required for oligomerization during the assembly of CBs, because mutant NTD^NLS^ is able to disrupt wild-type oligomers by interspersing coilin molecules that are unable to interact with other coilin molecules. Furthermore, the ability of R36A NTD^NLS^ to oligomerize allows the formation of hybrid oligomers with wild-type endogenous coilin. The results from R8A and D79A also strongly suggest that coilin-coilin interactions are mediated through more than one interface on the NTD. Therefore, an R8A and D79A double mutant NTD was tested for its ability to interact. Importantly, interactions with wild-type coilin were completely abolished by the double mutant (Figs 3e, S4b and S4c), showing that both R8 and D79 are necessary to form coilin oligomers and/or to associate with wild-type coilin molecules.

**Fig 5.**
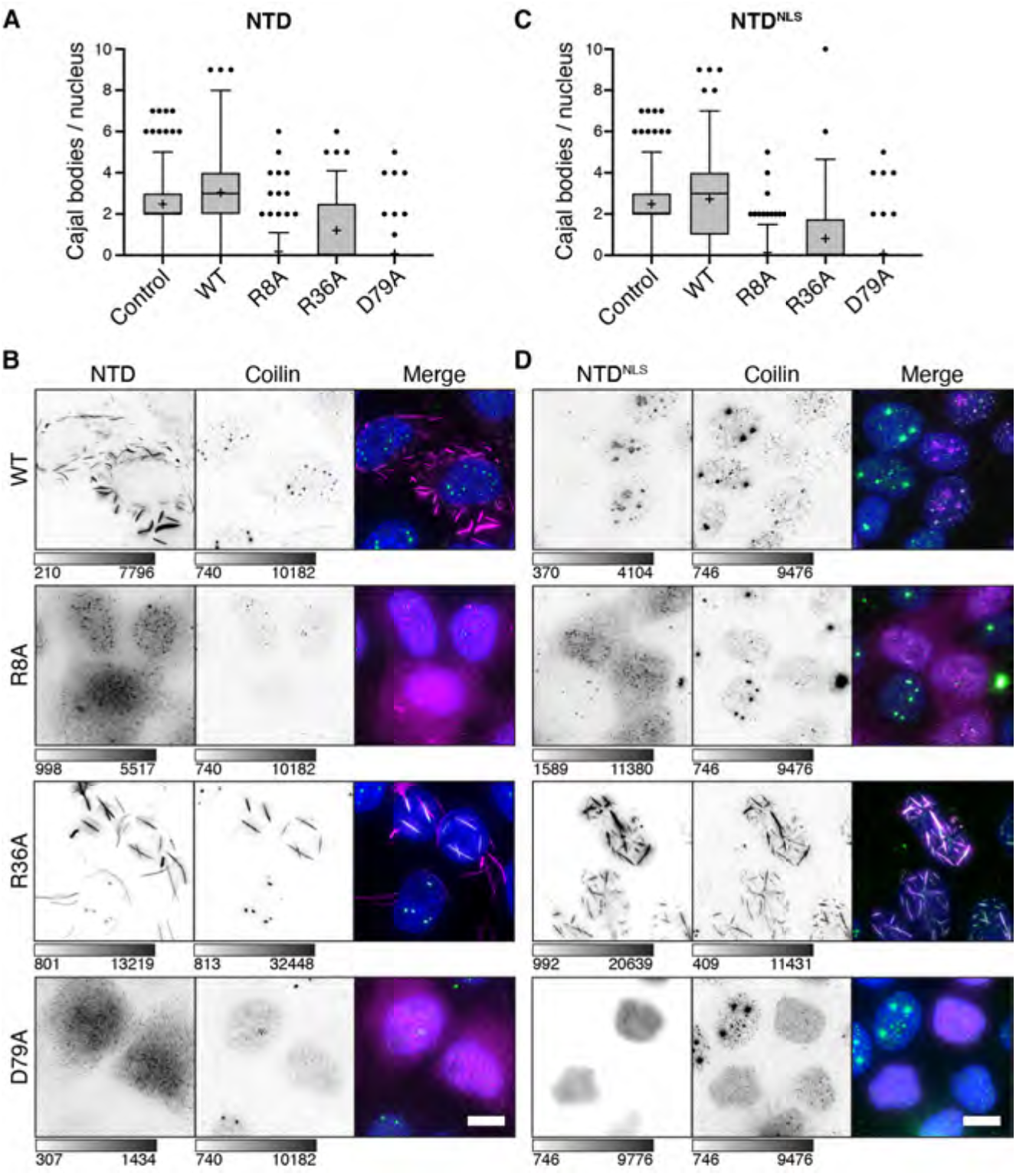
Coilin NTD mutants exert dominant negative effects on endogenous coilin and Cajal bodies. Expression of the myc-tagged coilin NTD in the cytoplasm of HeLa cells, immunostaining of NTD-myc and endogenous coilin, and counting of nuclear CBs is quantified in (A) and representative examples of mutant phenotypes are shown in (B), where immunostaining for myc is shown in magenta and dapi is shown in cyan. Identical analyses of the NTD^NLS^ constructs expressed in HeLa nuclei is shown (C&D). Horizontal line indicates the median, the cross indicates the mean. B&D) Grayscale bars given in analog-digital units. Scale bars = 10 μm. Images acquired with DeltaVision.

### Evidence that Nopp140 provides condensation properties to coilin oligomers

Based on these findings and on the predicted structures of both coilin^FL^ and its Nopp140 ligand (Fig. 6a), we hypothesized that Nopp140 must remodel coilin oligomers to yield an approximately spherical body in the nucleus. Although it is assumed that CBs are formed through biomolecular condensation, this has never been shown. One possibility is that coilin, which has an IDR linking the NTD with the structured CTD^31^, undergoes biomolecular condensation in addition to oligomerization through the NTD (Fig 6a). We tested this possibility, using the optodroplet assay^42^, which relies on light activation of the Cry2 dimerization domain to trigger condensation (Figs S5a and S5b) and which we previously used to implicate protein domains in biomolecular condensation^30^. Coilin^FL^ makes Cajal bodies without optical activation and additional puncta after induction, as expected (Fig. S5c). In contrast, coilin^ΔNTD^-Cry2 fails to form condensates in the absence or presence of light (Fig. 6B). Importantly, Nopp140 is comprised almost entirely of IDR and Nopp140^IDR^ readily forms optodroplets (Fig. 6c), leading to our model wherein the condensation properties of Nopp140 lead to remodeling of coilin oligomers and MLO formation. To test the most important prediction of this model – that Nopp140 is required to prevent coilin oligomerization in nuclei – we depleted Nopp140 from HeLa cells and expressed the coilin NTD^NLS^; as predicted, NTD^NLS^ oligomers formed in nucleoli (Fig. S5d), which was not observed when Nopp140 was present (see Fig. 1f). This suggests that coilin binding to Nopp140 modulates oligomerization by coilin, as depicted in Figure 6d.

**Fig 6.**
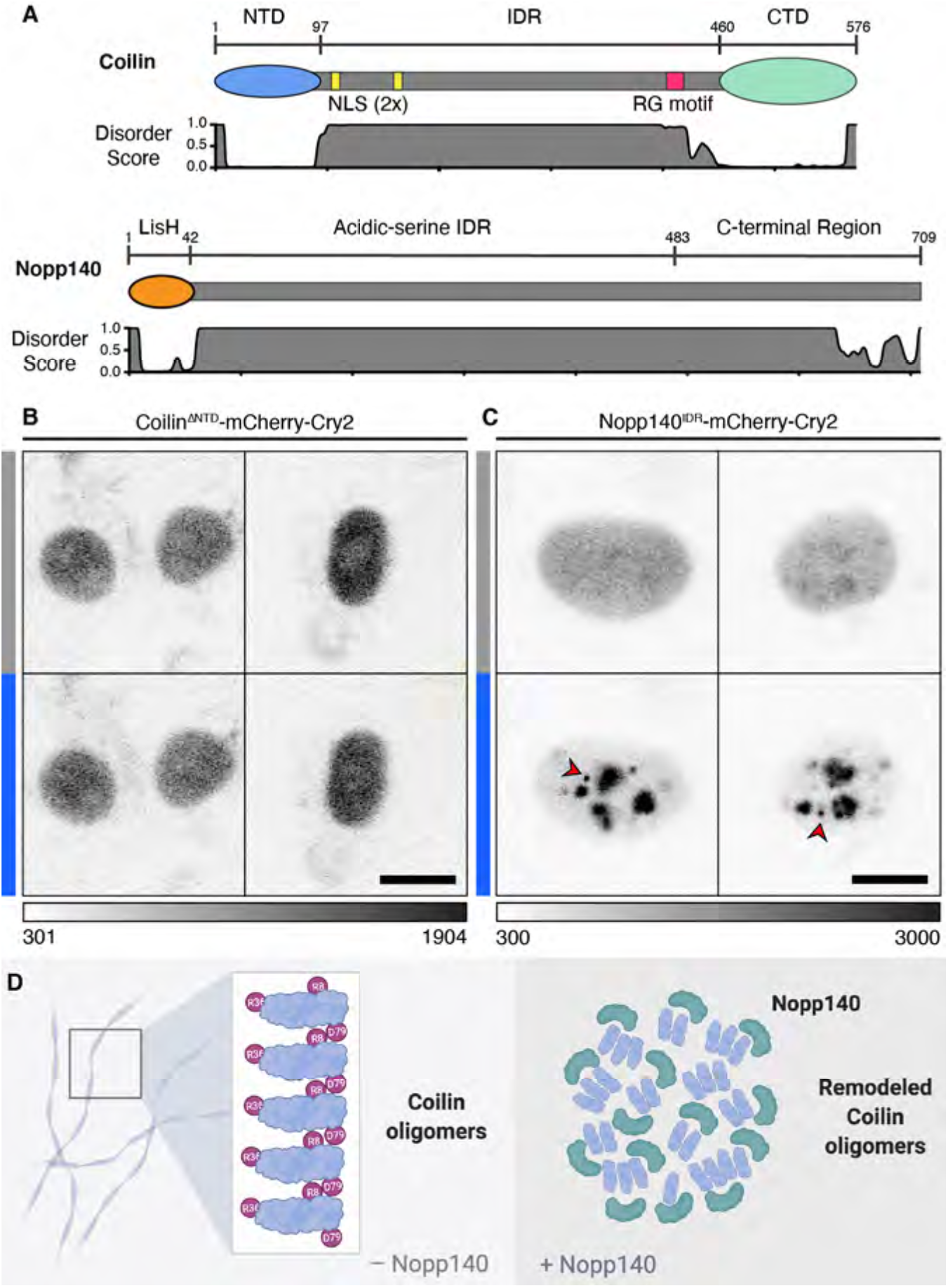
The Nopp140 IDR forms optodroplets, suggesting a mechanism for Cajal body assembly. A) Schematic representation of coilin and Nopp140 with overall domain structure and predicted secondary structure from RaptorX calculations. B) NIH-3T3 cells expressing Coilin^ΔNTD^ mCherry-Cry2 construct. Grayscale bar given in analog-digital units. Blue bars indicate light activation of the Cry2 domain (3 min). Scale bar = 10 μm. C) NIH-3T3 cells expressing Nopp140^IDR^ mCherry-Cry2 construct. Red arrowheads indicate non-nucleolar bodies formed by light induction. Scale bar = 10 μm. D) Schematic of proposed Nopp140 modification of coilin NTD oligomerization. Coilin NTD is drawn in blue and Nopp140 in green.

## Discussion

This study approaches the Cajal body as an endogenous cellular system, exploiting its conservation over millennia, to show the molecular details that drive its assembly and shape. The data enable us to transition from the knowledge that coilin is essential for the assembly and maintenance of CBs to a molecular model. We provide evidence that the coilin NTD is a folded domain with at least two distinct faces that mediate coilin-coilin oligomerization; coilin oligomers are remodeled by binding to Nopp140. *In vivo* FRET and co-immunoprecipitation data establish that the morphological effects of our single amino acid mutations are due to gain or loss of binding. For example, coilin binding to Nopp140 is lost when R36 is mutated to alanine, and coilin is observed as long, nuclear oligomers. Consistent with this and showing that the mutation does not induce an atypical behavior, wild-type coilin oligomerizes in the cytoplasm where Nopp140 is absent. Conversely, oligomerization is blocked when R8 or D79 are mutated, and neither oligomers nor nuclear bodies form. Taken together, the data indicate that *both* coilin oligomerization mediated by R8 and D79 *and* binding to Nopp140 mediated by R36 are required for CB assembly. Figure 3F shows a predicted structure suggesting that the coilin NTD contains two distinct interfaces that mediate inter-coilin interactions and confer oligomerization properties. Our working model (Fig. 6d) includes the inference that Nopp140 binds and modifies coilin oligomers through its ability to form condensates, either limiting their growth or changing the shape of the oligomers from linear to rounded.

Our findings highlight the importance of homo-oligomerization in the formation of healthy and diseased structures within cells^43–45^. Previous work has detected the formation of aggregates, oligomers, and fibrils in the context of droplet or hydrogel formation. For example, hnRNP A2 adopts a cross-β structure whether it is in hydrogels, droplets or in nuclei^46^. However, coilin oligomers do not appear to exhibit properties of amyloids, because they do not stain with Thioflavin T or Congo Red; moreover, they were negative for ProteoStat staining typical for protein aggregates (see Figs 1e and S1c). Oligomerization of other proteins, like NPM1 in nucleoli and SPOC in speckles, may underlie scaffolding or localization to the structure^47,48^. In our study, we found that the coilin NTD could not be replaced by a heterologous dimerization domain, indicating that higher-order oligomers are required for CB assembly. Moreover, coilin itself may not undergo biomolecular condensation *in vivo*, since coilin^ΔNTD^ was inactive in the optodroplet assay. Instead, our data and resulting model clarify an additional, essential role that Nopp140 plays in biomolecular condensation. First, we identified the NTD as the binding site for Nopp140 and furthermore validate its requirement in CB assembly by depleting Nopp140 and observing CB disruption. Second, we implicated biomolecular condensation by Nopp140 in the remodeling of coilin oligomers, by showing that Nopp140’s IDR is active in the optodroplet assay. This model suggests that coilin oligomers act with Nopp140 to form a minimal unit for assembly of the CB as we have seen with expression of the NTD^NLS^ protein in the cell nucleus (see Fig. 1c).

What, then, is the role of the rest of the coilin molecule? The RG box within the coilin IDR is modified by dimethylarginine required for SMN binding and the docking of gems to CBs^28,30^. The binding of Sm proteins to the CTD likely facilitates snRNP localization and maturation, linking CB assembly to its essential function^33^. The sites of interaction for additional coilin-binding proteins, such as the U6 snRNP component SART3 and the capping enzyme TGS1, as well as coilin-binding snRNAs and snoRNAs have not yet been mapped to particular domains on coilin^3,40^. Despite this incomplete information, the role of the coilin IDR could be simply to link the CB assembly function of the coilin NTD with the snRNP maturation function of the coilin CTD.

Our data and the resulting working model (Fig. 6d) invoke a process of coilin oligomerization that sheds new light on CB assembly. In 2005, Gall and colleagues speculated that CBs are “semi-fluid spheres suspended in semi-fluid nucleoplasm”^49^, representing possibly the first reference to liquid-liquid phase separation in the context of cellular compartments. This conclusion was based on observed shape, permeability, and differential protein concentrations between the nucleoplasm and the large CBs (10 μm in diameter) present in *Xenopus* germinal vesicles. Indeed, these coilin-containing “spheres” may be more closely related to histone locus bodies and are mostly extra-chromosomal^1,50,51^. Subsequent publications have referred to CBs as liquids^52^. However, electron microscopy shows what appear to be coiled electron dense fibers that led to the naming of “coiled bodies” before their identity with Cajal bodies was known^1,53^. Interestingly, FRAP data showed that, although 50% of coilin molecules recover within 1 minute of bleaching, the other 50% were immobile^11^. We speculate that this immobile coilin fraction is contained in longer-lived oligomers. Recent sub-diffraction fluorescence imaging has shown that CB morphology includes indentations or pockets to which other nuclear MLOs, such as gems, can be docked^30^. How coilin oligomers accommodate or facilitate these morphological relationships remains to be determined.

How do CBs achieve their dynamic character? We speculate that disassembly and reassembly – for example, before and after mitosis – must be mediated by the shortening of coilin oligomers or unbinding of Nopp140. Although there is *a priori* no reason to suspect that oligomerization of the coilin NTD would be altered by cell cycle, coilin is phosphorylated and methylated at several sites suggesting that post-translational modifications of coilin and its binding partners might contribute to these changes^34,54^. Moreover, it seems likely that the increase in Nopp140 solubility during mitosis, which is caused by extensive phosphorylation by cdc2 kinase, may play a role in CB dynamics^55,56^. Nopp140 is a shared component between CBs and nucleoli, and nucleoli also disassemble at mitosis. Inevitably, these changes must also be linked to the dependency of nucleoli and CBs on transcription, which nucleates these MLOs at active rDNA and snRNA genes, respectively^9,44,57^. Thus, the role(s) of transcription, RNA, post-translational modifications and additional protein binding partners are leading towards a molecular model for CB dynamics. Importantly, the discovery that oligomerization by coilin and binding to Nopp140 are required for the limited assembly of an approximately spherical membraneless structure suggests this may be a core CB particle that provides a basis for recruitment of the full complexity of CB components.

## Acknowledgments

The authors wish to thank members of the Neugebauer lab for helpful discussions and comments on the manuscript. We thank Greg Matera for the *Coil-/-* MEF cell line. The study was supported by NIH awards NINDS-F31NS105379 (to E.M.C) and U01CA200147 TCPA-2017-Neugebauer (to K.M.N.). J.E. was supported by the Yale BioMed Amgen Scholars Program through a grant from the Amgen Foundation (41716881). This work is solely the responsibility of the authors and does not necessarily represent the official views of the NIH.

## Author contributions

M.M., E.M.C. and K.M.N. designed the study. M.M., S.S. and K.S. generated all constructs. M.M., E.M.C., S.S., J.E., and K.S. prepared cell lines and performed experiments. E.M.C. and J.E. carried out image analysis. K.M.N. supervised the study. All authors contributed to writing the manuscript.

## Methods

### Experimental Model

#### Tissue Culture Cell Lines

The *coil^−/−^* Mouse Embryonic Fibroblasts were a gift from Greg Matera^15^. HeLa cells are from the Kyoto linage (RRID: CVCL_1922). Cry2 optodroplet experiments were performed in NIH-3T3 cells obtained from ATCC (# CRL-1658). 293FT cells were used as a lentivirus packaging line only and were obtained from Thermo (#R70007). All cell lines were cultured in DMEM supplemented with 10% FBS, penicillin, and streptomycin (Gibco) at 37°C in a 5% CO_2_ atmosphere. Cells were regularly screened for the presence of mycoplasma infection.

### Method Details

#### Plasmid and Viral Vector Construction

Coilin and Nopp140 constructs were cloned into the pEGFP-N1 backbone modified to include a C-terminal c-myc or HA epitope tag, rather than a fluorescent protein. Coilin and Nopp140 constructs used for FRET assays were cloned into pECFP-N1, pECFP-C1, pEYFP-N1, or pEYFP-C1 depending on the measurement. Cry2 constructs were inserted into the pHR_SFFV backbone originally received as a gift from Cliff Brangwynne^42^. Lentiviral particles were generated by co-transfecting the pHR_SFFV vector with pCMV 8.74 (generated by Didier Trono, Addgene #22036) and pMD2.G (generated by Didier Trono, Addgene #12259) into 293FT cells using Fugene HD transfection reagent (Promega). Viral particles were harvested by removing supernatant and filtering through 0.45μm syringe filters to remove debris.

#### Mammalian Cell Expression and Imaging Sample Preparation

Transient transfections were performed using Lipofectamine 3000 reagent (Invitrogen) in cells grown on No. 1.5 coverslips (Zeiss). Cells were fixed in fresh 4% paraformaldehyde (Sigma) 24 hours post-transfection. Each sample was then blocked and permeabilized in 3% Bovine Serum Albumin (Sigma) and 0.1% Triton X-100 (American Bioanalytical). Immunofluorescence labeling was performed in the same blocking buffer using sequential primary antibody followed by secondary antibody labeled with fluorophore. Nuclei were stained with 1 μg/mL Hoechst 33342 (Invitrogen). Coverslips were then mounted using DABCO

For live-cell imaging of Cry2 constructs, stable NIH-3T3 cell lines were generated by applying virus to wild-type cells. Cells were then plated onto glass bottomed 35 mm dishes (MatTek). The media was replaced with Live Cell Imaging Solution supplemented with 20 mM glucose and 1 μg/mL Hoechst.

For FRET assays, cells were transfected with pairs of CFP and YFP constructs using the conditions described above, fixed in 2% paraformaldehyde and mounted in DABCO without Hoechst.

#### siRNA Silencing

Coilin and Nopp140 were depleted using pools of Ambion Silencer Select oligos (ThermoFisher). siRNAs targeting *Coil* (s15662, s15663, s15664) and *Nolc1* (s17632, s17633, s17634) were prepared according to the manufacturer’s protocols and combined into pools of equimolar amounts of each of the three oligos. A total of 10 pmol were transfected into HeLa cells grown to 50% confluency in a 12-well plate with Lipofectamine RNAiMAX (Invitrogen). After 24 hours, cells were re-plated onto coverslips for imaging or for western blot analysis and harvested or fixed after 48 hours post-transfection.

#### Microscopy Platforms

Fixed cell samples were imaged on one of two platforms as noted in the figure legends. For those imaged using the DeltaVision platform (Applied Precision), a 60x PlanApo N.A. 1.42 Oil objective (Olympus) was used to image the full depth of the sample at 0.2 μm intervals. The resulting wide field image stacks were deconvolved with the native Applied Precision software.

For samples imaged using laser scanning confocal, a Leica Sp8 was used with a 63x HC PL APO CS2 Leica objective. The instrument is equipped with a white light laser, three hybrid detectors and two PMT detectors for spectral selection, and channels were imaged sequentially to eliminate bleed through.

Live-cell imaging of Cry2 samples was performed on a Bruker Opterra II Swept Field Instrument. Samples were simultaneously illuminated with 488 nm and 561 nm laser light, where the 488 nm light activates Cry2 and the sample was imaged in the 561 nm channel to detect mCherry. A PlanApo 60X 1.2 NA water immersion objective was used and the instrument is equipped with a Evolve 512 Delta EMCCD camera. Ten frames at one second each were captured before activation for a total of 180 seconds. A stage-top incubator was used to maintain cells at 37° throughout the imaging protocol.

#### FRET Measurements

Förster resonant energy transfer (FRET) measurements were made using the acceptor photobleaching scheme^58^. Measurements were made on a Zeiss 710 laser scanning confocal instrument equipped with 458 nm and 514nm lasers for imaging CFP and YFP respectively, detected by corresponding spectrally selective PMT detectors. Measurements were made by imaging both channels in a given region of interest (ROI) followed by a high intensity bleaching scan of the ROI with the 514nm laser. The same ROI was then reimaged in both channels. A fusion construct of CFP-YFP was used as a positive control and freely expressed non-fused CFP and YFP were used as a negative control.

#### Immunoprecipitation Assays and Western Blotting

Immunoprecipitation assays were performed by transfecting a bait GFP fusion protein into HeLa cells as discussed above. Approximately eight million cells were harvested for each sample and lysed into Pierce lysis buffer (Thermo) containing Complete protease inhibitors (Roche). Lysate was clarified by centrifugation at 10,000 g for 30 min. Pellet and a portion of supernatant were reserved as input. The remaining lysate was incubated with 4 μg of goat anti-GFP overnight, followed by immunoprecipitation with Protein G Magnetic SureBeads (BioRad). Beads were washed three times in Pierce lysis buffer and eluted into NuPAGE Sample Buffer (Invitrogen). The resulting eluent was run onto a NuPAGE 4-12%, Bis-Tris gel (Thermo), transferred to nitrocellulose and probed with primary antibody to be visualized with horseradish peroxidase conjugated secondary antibody.

#### Amyloid Dye Staining

Amyloid dye samples were prepared for microscopy as described above for immune fluorescence. Following antibody probing with Cy5 to prevent bleed through, samples were stained with either Thioflavin T (200 nM, Sigma), Congo Red (0.005%, Sigma), or Proteostat (3 μM, Biotium) solutions.

### Quantification and Statistical Analysis

#### FRET Analysis

The bleached portion of each image was cropped to a thirty-by-thirty pixel ROI. FRET efficiency percentage for that ROI was calculated as E_f_ = (I_2_ − I_1_) / (I_2_) where I_2_ is the mean intensity of the CFP channel after photobleaching of YFP, and I_1_ is the intensity of the CFP channel before photobleaching. I_1_ and I_2_ were background corrected by subtracting the mean intensity of five ROIs outside of cells. The reported FRET efficiency is the mean of twelve ROIs from two separate biological replicates with error bars as standard error of the mean.

#### Nuclear Body and Coilin Mutant Phenotype Quantification

Where phenotypes or numbers of nuclear bodies are quantified, cells were counted by hand. At least two biological replicates per condition were analyzed. An author previously unfamiliar with the data was asked to establish phenotype categories and count the number of positively transfected cells matching those categories. Likewise, they counted the number of nuclear bodies per nucleus in conditions where nuclear bodies were quantified. The sample size is noted in the respective figure legends.

#### Software

Images were prepared for figures using ImageJ (FIJI) version 2.0.0-rc-69/1.52p^59^. Structure prediction was performed on the Jpred and RaptorX server using default parameters^41,60^. Coilin alignments were performed using Clustal Omega using the default parameters^61^.

#### Data and code availability

This study did not generate large datasets or computer code.

---

**Figure S1.**
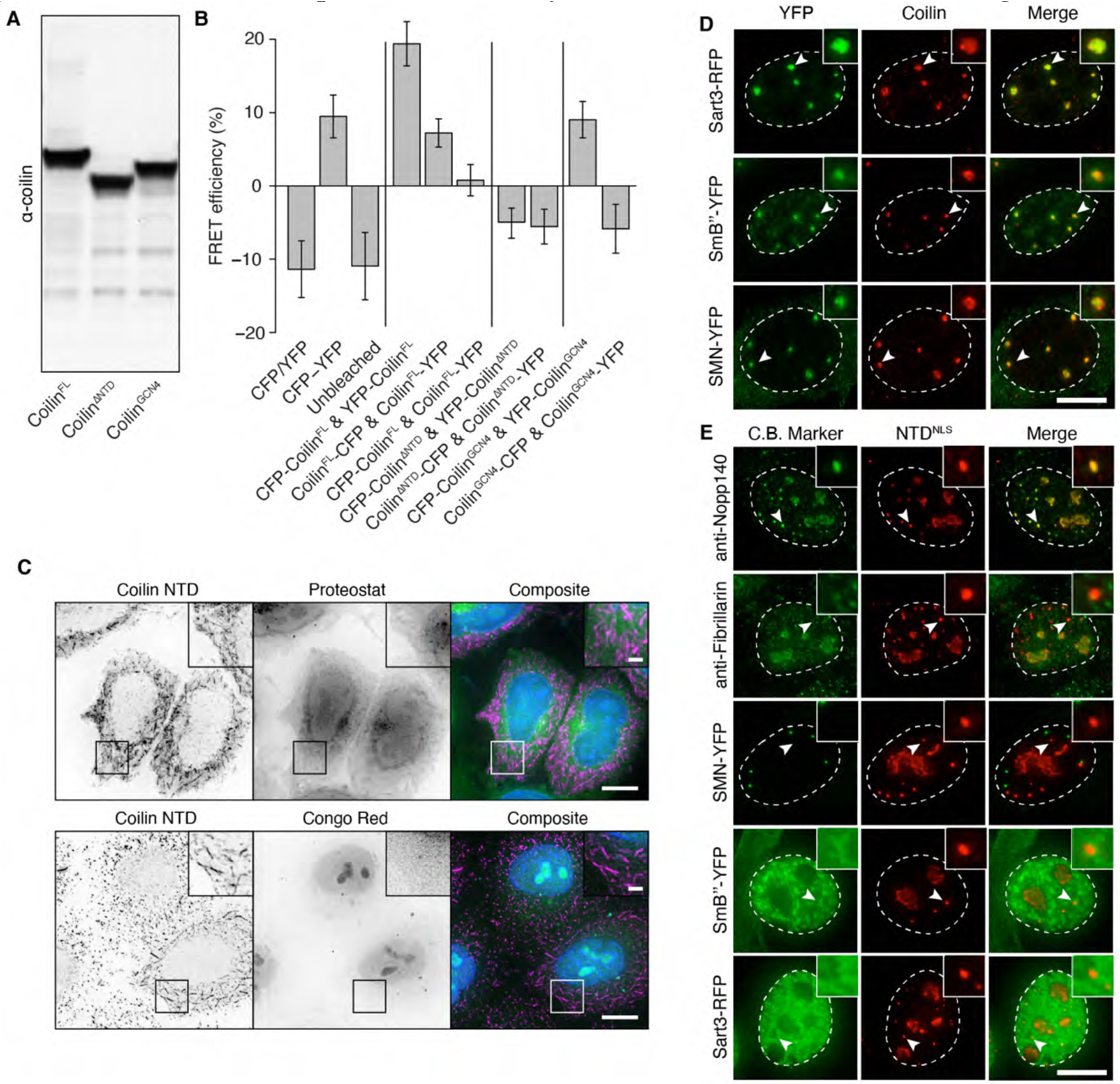
The coilin NTD promotes non-amyloid self-interaction; related to Figure 1. A) Western blot of lysate from *coil ^−/−^*MEFs expressing transiently transfected coilin constructs. B) FRET analysis performed in *coil ^−/−^*MEFs with fluorescent protein donors and acceptors included at both N and C termini. Negative values are due to photobleaching of the CFP donor. Error bars indicate standard deviation, n = 12. C) NTD expressed in HeLa and counterstained with dyes sensitive to amyloid cross-β structures. Scale bar = 10 μm, Inset scale bar = 2 μm. D) YFP-tagged Cajal body markers co-transfected with Coilin^FL^ into *coil ^−/−^*MEFs. Arrowheads indicate example Cajal bodies enlarged in the inset. Scale bar = 10 μm. E) Primary antibody staining or transfection of YFP-tagged Cajal body markers in *coil ^−/−^*MEFs transfected with NTD^NLS^. Arrowheads indicate NTD bodies enlarged in the inset. Scale bar = 10 μm. All images acquired with DeltaVision.

**Figure S2.**
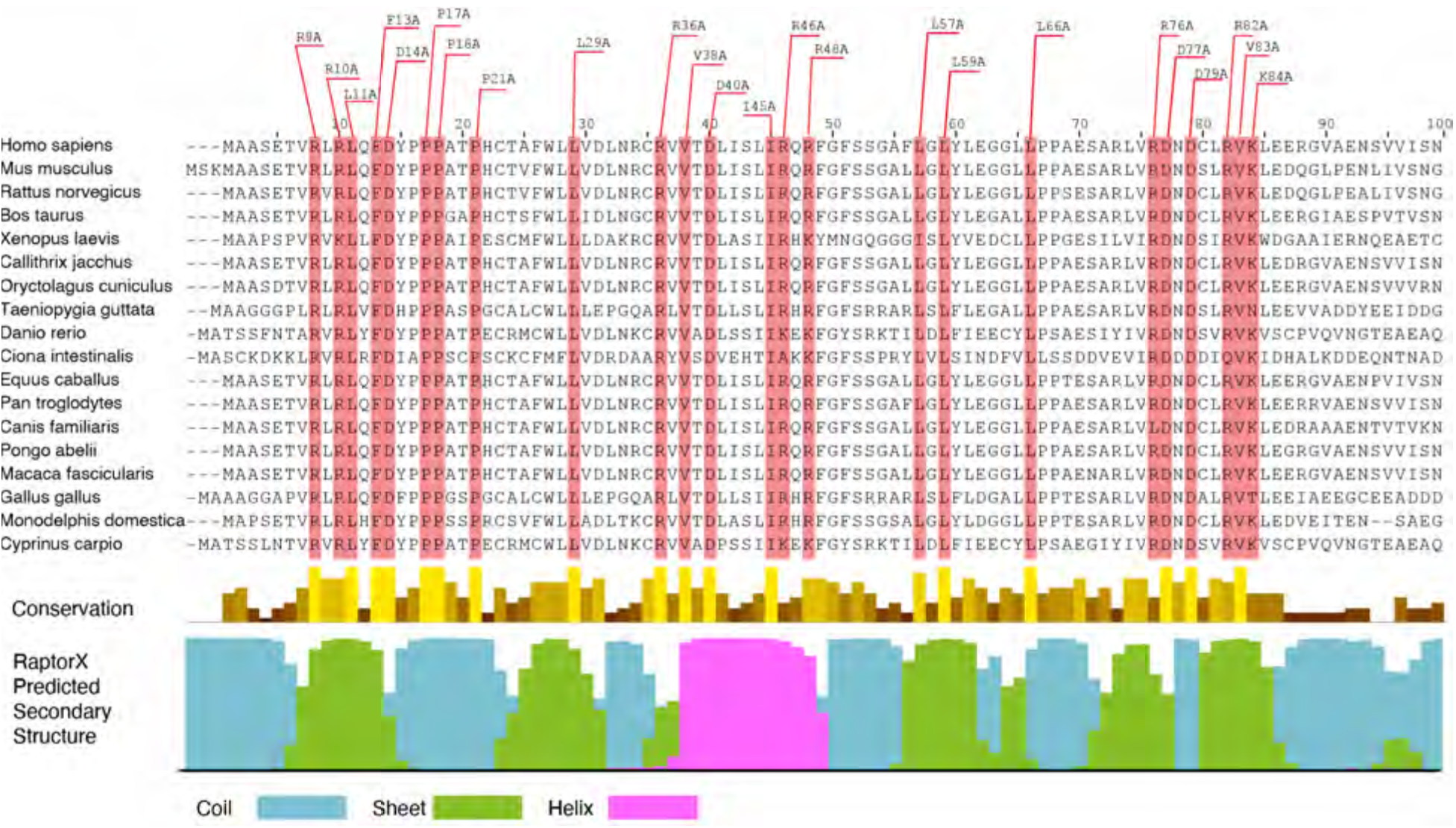
Alignment of vertebrate coilin sequences indicates highly conserved residues; related to Figure 3. Multiple sequence alignment of vertebrate coilin orthologs, cropped to the first one hundred amino acids. Conservation track is computed in Clustal Omega and reveals highly conserved residues selected for alanine scanning mutagenesis, highlighted in red. RaptorX secondary structure predictions indicate possible regions of beta-strands (green) or alpha-helix (magenta).

**Figure S3.**
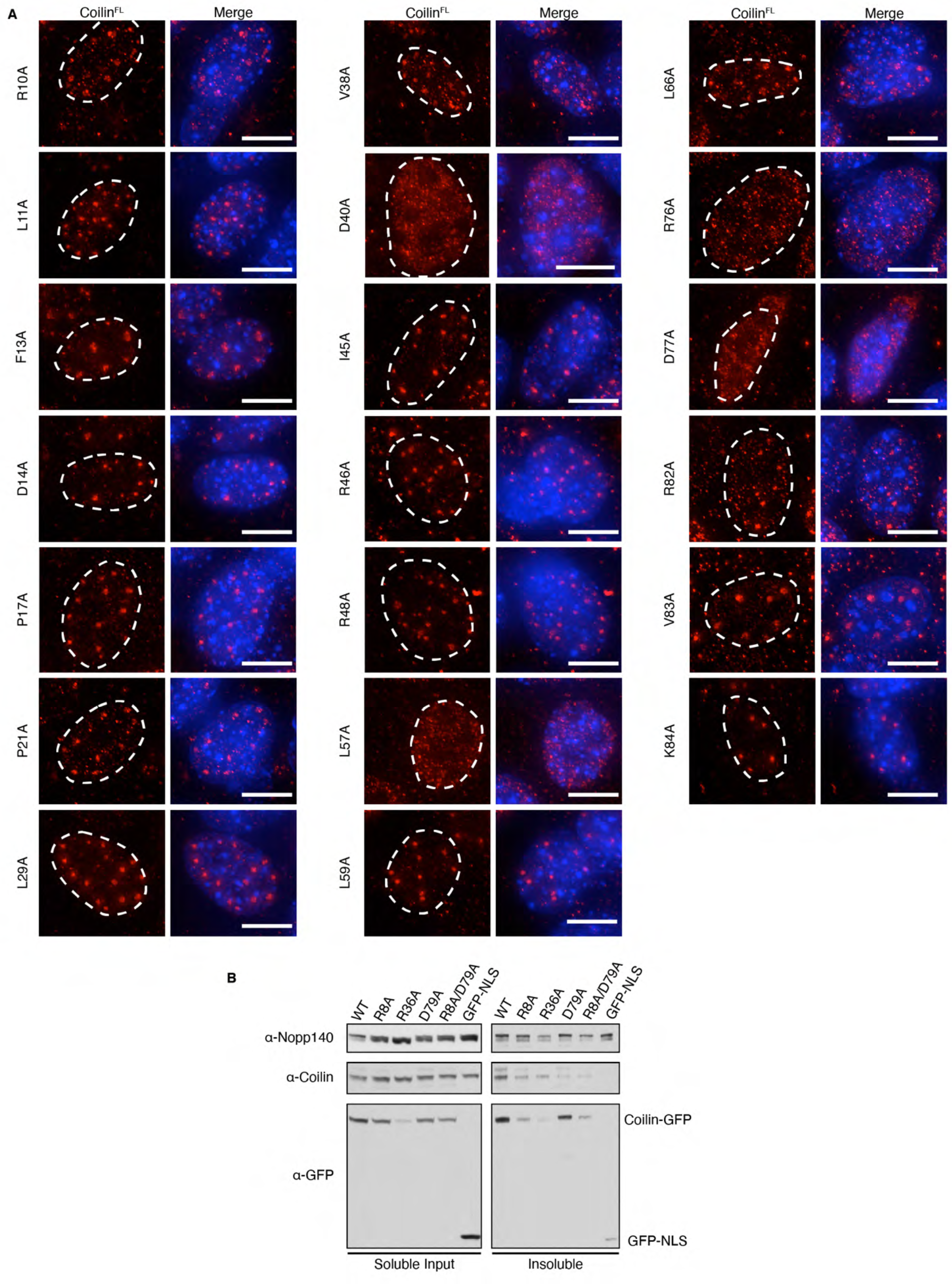
Mutation of conserved residues reveals key interaction sites of the coilin NTD; related to Figure 3. A) Coilin transfected into *coil ^−/−^*MEFs and visualized by immunofluorescence. Mutations correspond to highlighted residues in Figure S2, with numbering according to the human sequence. Blue indicates nuclear staining. Images acquired with Delta Vision. Scale bars = 10 μm. B) Soluble Input and insoluble pellet from immunoprecipitation shown in Figure 3. Lysates were produced from HeLa cells.

**Figure S4.**
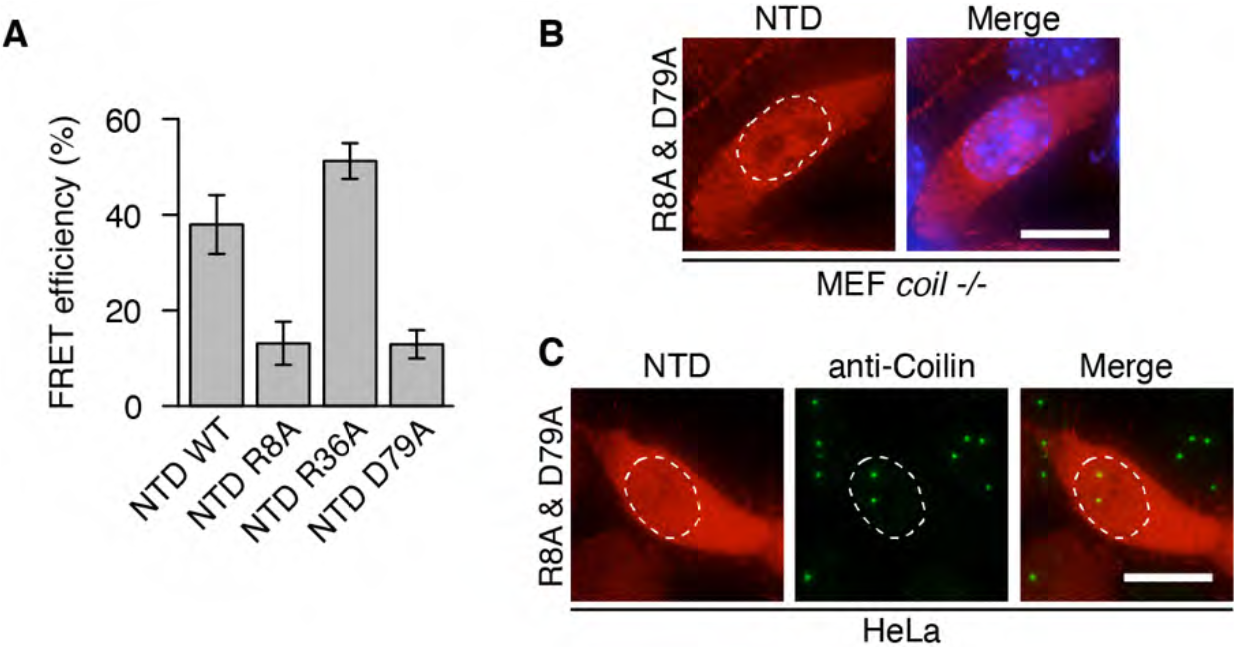
Point mutations of the NTD disrupt its oligomerization; related to Figures 4 & 5. A) FRET analysis of CFP and YFP labeled coilin NTD are co-transfected into *coil ^−/−^*MEFs. Error bars indicate standard deviation. B) Myc-tagged coilin NTD bearing both R8A and D79A mutations transfected into *coil ^−/−^*MEFs. Scale bar = 10 μm. C) Coilin NTD bearing both R8A and D79A mutations transfected into HeLa. In B and C, cells are immunostained for the myc-tag (red) and, in C, endogenous coilin (green). Images acquired with DeltaVision. Scale bar = 10 μm.

**Figure S5.**
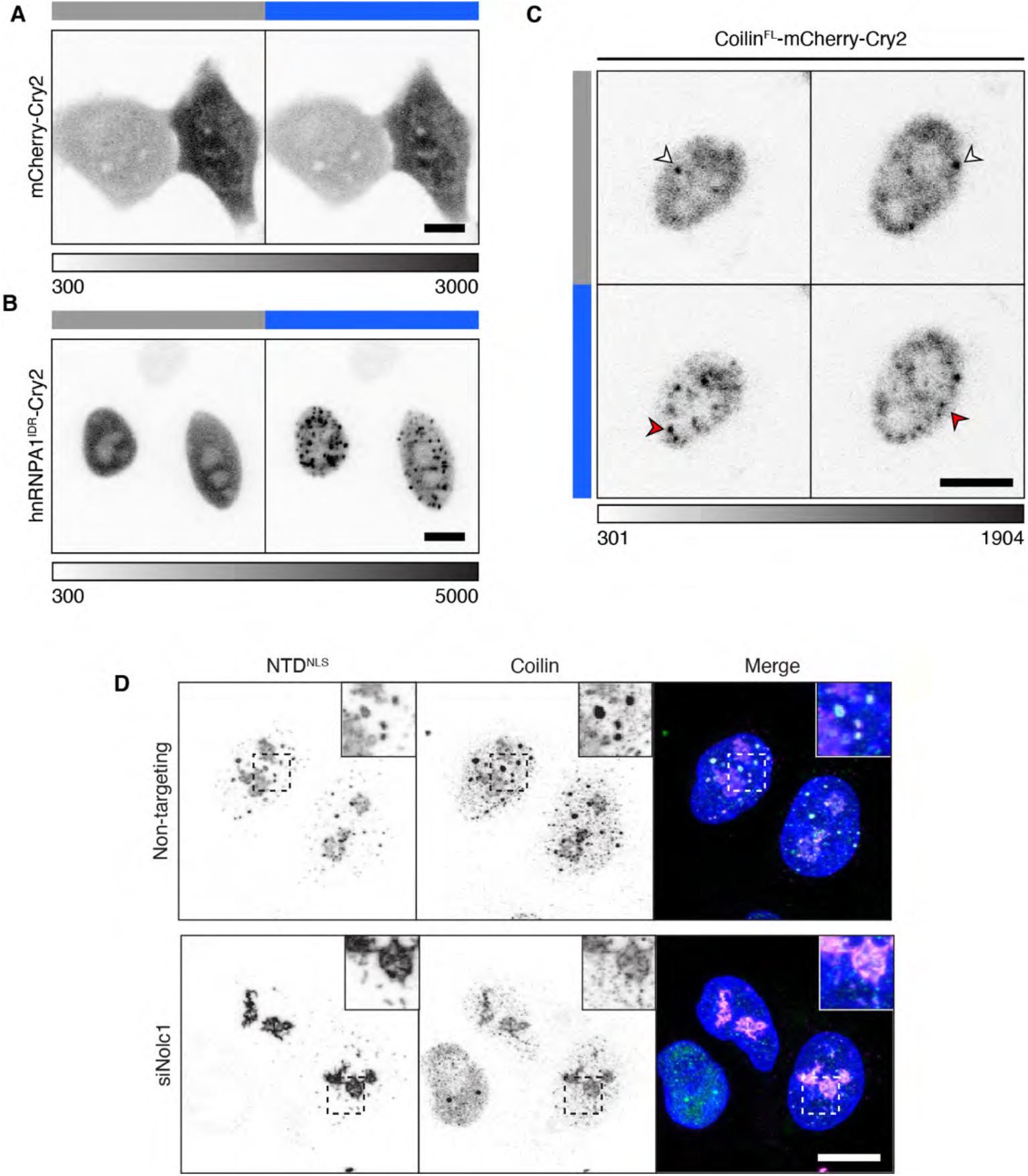
Evidence that Nopp140 drives Cajal body condensation; related to Figure 6. A&B) Live-cell imaging of NIH-3T3 cells expressing mCherry-Cry2 (A) and the hnRNP-A1 intrinsically disordered region (B). Blue bar indicates 180 s activation with blue light. C) Coilin^FL^-mCherry-Cry2 forming Cajal bodies before activation of Cry2 by blue light (white arrows) and additional bodies formed upon 3 min of activation (red arrows). D) HeLa cells co-transfected with siRNA and the coilin NTD^NLS^, and counterstained for coilin, imaged with laser scanning confocal microscopy. Grayscale bars given in analog-digital units. Scale bars = 10 μm.

